# A model of task-level human stepping regulation yields semistable walking

**DOI:** 10.1101/2024.03.05.583616

**Authors:** Navendu S. Patil, Jonathan B. Dingwell, Joseph P. Cusumano

**Affiliations:** Department of Kinesiology, Pennsylvania State University, University Park, PA 16802, USA; Department of Engineering Science & Mechanics, Pennsylvania State University, University Park, PA 16802, USA

## Abstract

A simple lateral dynamic walker, with swing leg dynamics and three adjustable input parameters, is used to study how motor regulation affects frontal plane stepping. Motivated by experimental observations and phenomenological models, we imposed task-level multiobjective regulation targeting the walker’s optimal lateral foot placement at each step. The regulator prioritizes achieving step width and lateral body position goals to varying degrees by choosing a mixture parameter. Our model thus integrates a lateral *mechanical template*, which captures fundamental mechanics of frontal-plane walking, with a lateral *motor regulation template*, an empirically verified model of how humans manipulate lateral foot placements in a goal-directed manner. The model captures experimentally observed stepping fluctuation statistics and demonstrates how linear empirical models of stepping dynamics can emerge from first-principles nonlinear mechanics. We find that task-level regulation gives rise to a goal equivalent manifold in the system’s extended state space of mechanical states and inputs, a subset of which contains a continuum of period-1 gaits forming a *semistable* set: perturbations off of any of its gaits result in transients that return to the set, though typically to different gaits.

## 1 Introduction

Humans are highly versatile bipedal walkers. Having learned as toddlers to take steps without falling, people can walk purposefully in a myriad of real-world situations. Successful walking requires, among other things, realizing an appropriate sequence of footsteps [1]. While walking, humans can perform goal-directed tasks, like staying on a path or walking alongside another person, with equal success via any one of infinitely many possible sequences of footsteps. Such task-level equifinality of foot placements, along with body-level biomechanical redundancies in muscles and joints [2], physiological motor noise [3], and environmental disturbances, all contribute to the significant variability observed in human walking [4–6]. Indeed, step-to-step observations of human walking provide a window into how the nervous system manages redundancies and equifinality while achieving task goals in specific walking contexts. However, the non-uniqueness of solutions to any given walking task makes the modeling and analysis of such systems mathematically ill-posed.

We previously posited that humans achieve stable walking via a hierarchical schema consisting of two functionally distinct sets of processes that we distinguish as *control* versus *regulation* [7]: whereas “control” refers to processes required for a walker to remain *viable*, i.e., to take steps without falling, “regulation” designates step-to-step processes a viable walker needs to carry out specific goal-directed *tasks*. The notion of viability [8] is well-suited to handle the ill-posedness of walking as it merely characterizes the walker’s ability to avoid falls and is, therefore, agnostic to the existence of attracting steady states or goal-directedness. Conversely, by itself, regulation at the task level does not aim to ensure the walker’s viability: on the contrary, it assumes viability as a prerequisite. The overall control architecture that results mirrors those obtained in engineered systems using *successive loop closure* [9, 10] with, in our case, the “control” subsystem responsible for viable stepping acting as the “inner loop” and the goal-directed “regulator” subsystem as the “outer loop”. Moreover, its hierarchical structure is consistent with the neurophysiological architecture of perception-action processes responsible for human motor performance [11, 12].

People, particularly older adults, frequently fall while walking, and can experience serious injuries, such as hip fracture, especially after falling sideways [13, 14]. Taking wider steps can modify the lateral motion of the body’s center of mass (CoM) to prevent such falls [1, 15, 16]. Thus, to reduce the incidence of falls, it is imperative to understand the dynamics of step-to-step foot placements in human walking, especially in the lateral direction. In previous experimental work [4–6], we used simple linear update models based on error minimization with respect to hypothesized walking task goals, or *motor regulation templates*, to study the variability structure of human stepping time series. Multiobjective models that strongly prioritized regulating step width over lateral body/CoM position replicated the fluctuation statistics of human lateral foot placements (figure 1). However, given that these empirical models are developed using only task-level stepping data, without considering underlying biomechanics, their effectiveness is, perhaps, surprising. It remains an open question how the Newtonian mechanics of the body, which is essentially nonlinear, can give rise to such linear regulation at the task level.

**Figure 1.**
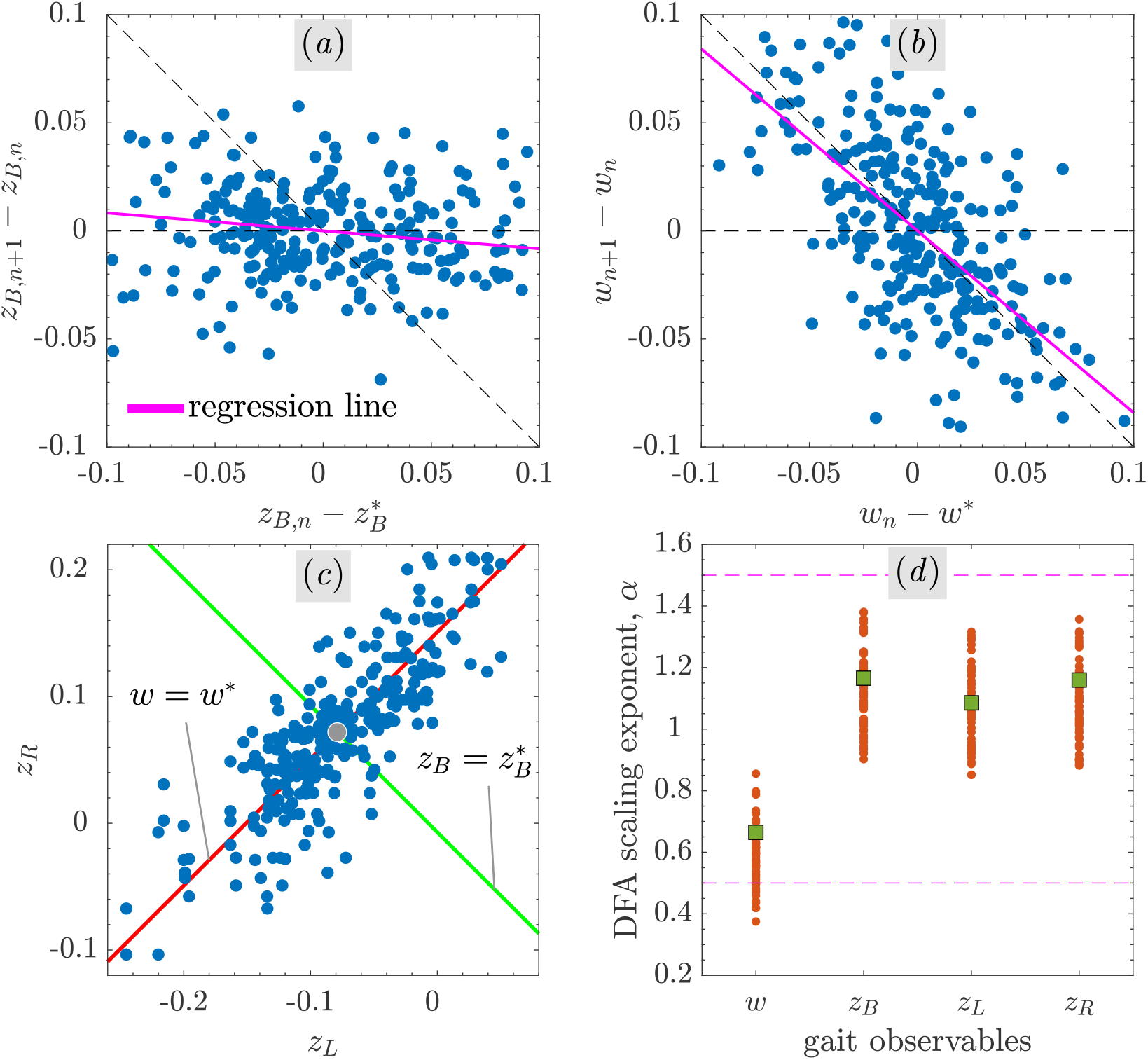
Human walking data showing step-to-step multiobjective regulation of lateral body position, *z*_*B*_, and step width, *w* [6]. Typical experimental “direct control” plots [6] in (*a*) body position and (*b*) step width show how deviations from their respective target values (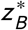 and *w* ^∗^) are corrected from one step to the next. Goals 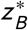 and *w* ^∗^ are set to experimental averages 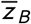 and 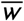, respectively. Linear regression shows that fluctuations in *z*_*B*_ are largely uncorrected (*regression line* slope ≈ 0); those in *w* are strongly corrected (slope ≈ −1). Same walking trial plotted in the stepping plane (*c*) of left and right foot placements (*z*_*L*_, *z*_*R*_) shows much larger fluctuations in *z*_*B*_ than *w*, where 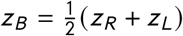 and *w* = *z*_*R*_ − *z*_*L*_. Only point at intersection (*filled circle*) of constant body position 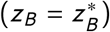 and constant step width (*w* = *w* ^∗^) goal equivalent manifolds (GEMs) perfectly satisfies both goals. Detrended fluctuation analysis (DFA) exponents, *α*, for different stepping observables (*d*): a fluctuation at step *n* is likely to be followed by a fluctuation at step *n* + 1 of the same sign (i.e., it is persistent) if *α* > 0.5; of opposite sign (i.e., antipersistent) if 0 < *α* < 0.5; *α* = 0.5 for white noise, while *α* = 1.5 for a pure random walk (*horizontal broken lines*). DFA *α* from a pooled data set (*dots*: 65 treadmill walking trials of 290 steps each; *filled squares*: sample trial in *a*-*c*) show much lower degree of statistical persistence in step width fluctuations than in body position and foot placements, consistent with the stronger regulation of *w*. Linear stochastic update models, or motor regulation templates, capture these statistical features of step-to-step fluctuations of observables {*z*_*B*_, *w*} (or, {*z*_*L*_, *z*_*R*_}). See [6] for detailed study.

To address this question, we here couple a mechanical template [17, 18] for frontal plane stepping on level ground with an empirically verified motor regulation template [6]. The resulting *integrated template model* shows how motor regulation behavior observed in humans can indeed be obtained from mechanical first principles. We demonstrate that the apparent decoupling between dynamics at the task and body levels observed in experiments arises naturally from the hierarchical regulation scheme “embodied” in our model. Furthermore, we show how task-level optimal regulation creates a goal equivalent manifold (GEM) [19] at the “body-level”, i.e., in the system’s extended state space of mechanical states and inputs.

Steady walking has commonly been thought of as a limit cycle, i.e., an asymptotically stable periodic gait [20, 21], possibly perturbed by noise [22, 23]. To remain versatile, however, humans have necessarily learned to walk with many periodic gaits, with different observables such as step length and step time, both stably and efficiently. Here, we find a manifold in the extended state space of the walker that is filled with periodic gaits, each of which is capable of achieving task-level goals. Furthermore, we show that at least a portion of this manifold forms a *semistable set* [24–27], i.e., a continuum of nonisolated periodic gaits such that perturbations off of any one of them result in transient dynamics that approach other, *typically different*, periodic gaits in the set. The semistability of walking, demonstrated here for the first time, is a direct consequence of equifinality: it is less constraining than the asymptotic stability of limit cycles and is, therefore, more suitable to characterize stability of versatile biological walkers like humans. Moreover, semistability is consistent with the idea that humans can, at least in principle, remain viable (i.e., avoid falls) [8] by switching between regulation strategies and their corresponding steady-state periodic gaits [7].

## 2 Task-level regulation in the frontal plane

We adapt a 2D compass walker originally used to model sagittal-plane walking [18, 28] to study powered dynamics in the *frontal plane* on level ground (figure 2). The walker can adjust three input parameters {*P*, *k*, 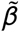} at the beginning of each walking step: the impulsive push-off input *P* ⩾ 0 models ankle actuation during toe-off in humans; a linear torsional spring, with stiffness *k* > 0 and unstretched angle 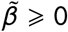, models hip actuation during the stance phase [18, 29].

**Figure 2.**
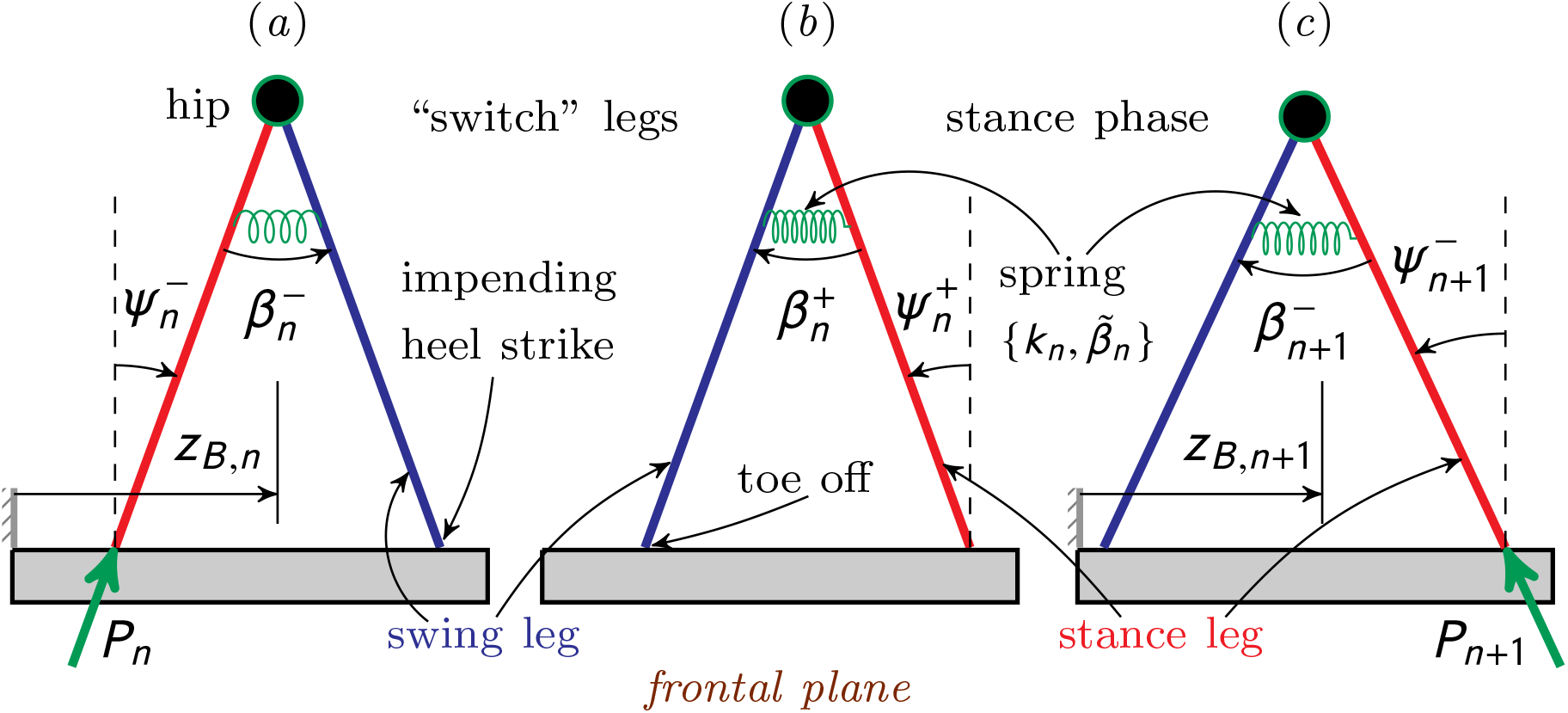
Three snapshots of a 2D powered compass walker with hip spring [18] adapted to walk laterally in the *frontal plane* on level ground: (*a*) just before *n*^th^, (*b*) just after *n*^th^, and (*c*) just before (*n* + 1)^st^ heel strike. The walker rocks back-and-forth in the frontal plane with anteroposterior axis directed into/out of the page. The walker has straight, massless, stance (*red*) and swing (*blue*) legs, and a point mass *M* at the hip (*circle*); point masses, *m*, at feet are infinitesimally small compared to *M*. Push-off impulse, *P*, is applied instantaneously just before heel strike (at which 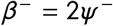) at the end of stance phase. Additional input parameters: torsional spring (*coil*) at hip, stiffness *k* and unstretched angle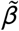, can be adjusted instantaneously at beginning of each stance phase. The walker’s mechanical state in the lab frame just before applying *P*_*n*_ is 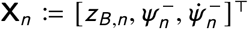 (see text).

### 2.1 Discrete-time stepping dynamics

Every walking step begins with an instantaneous impulsive double-support phase (figure 2*a*-*b*) followed by a continuous-time single-support stance phase (figure 2*b*-*c*) so the walker’s step-to-step dynamics are inherently *hybrid* [30].

The stance leg’s mechanical state just before applying push-off *P*_*n*_ is 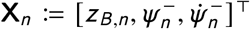, where *z*_*B*_ is the lateral body position in the inertial lab frame, *ψ* ^−^ is the stance leg angle in the frontal plane, and 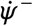 is its angular rate. The overall dynamics of the instantaneous double-support phase, consisting of push off, followed by a heel strike, and “switching” of stance and swing legs, is governed by the *impact map*:

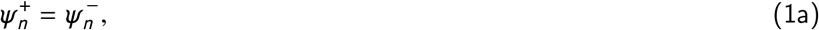

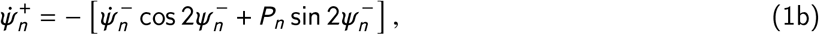

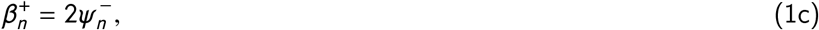

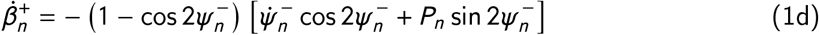

where the superscripts − and + indicate instants just before push off and just after heel strike, respectively. The walker’s state just after *n*^th^ heel strike, 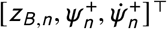, is independent of its swing leg angular rate 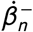 just before push off (equation 1) [28]. Consequently, the stance leg’s state **X**_*n*_ is, in fact, also the walker’s state necessary to evolve its step-to-step dynamics forward in time.

During the *n*^th^ continuous stance phase (figure 2), the walker’s dynamics is governed by the ordinary differential equations (ODEs):

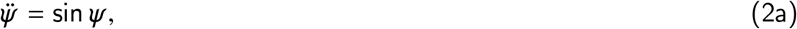

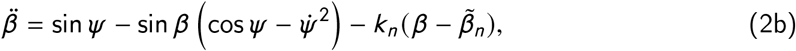

beginning from a state just after heel strike that lies on a codimension-two manifold 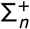 in the above ODEs’ 4D state space, and evolves to a state just before next push off that lies on a codimension-one Poincaré section 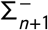, where

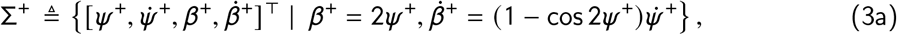

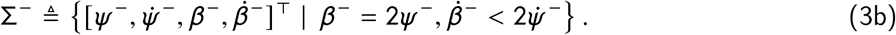

The spring parameters 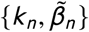 remain fixed during the *n*^th^ stance phase and do not affect the walker’s discrete dynamics (equation 1) for massless legs with infinitesimally small feet (figure 2). The above equations of motion (equations 1-3) are expressed in nondimensionalized variables, viz., time 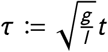, impulse 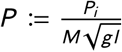, stiffness 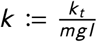, where *t*, *P*_*i*_, *k*_*t*_ are their dimensional counterparts, respectively, along with gravity *g*, leg length *l*, hip mass *M*, and foot mass *m* (figure 2).

Walking motion is feasible when 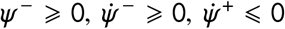 The stance leg does not swing past vertical if

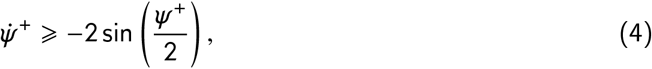

thus precluding crossover steps. Consequently, we get an inequality constraint on the states and inputs (using equations 1 and 4):

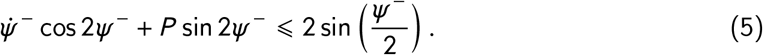

Using equation (1) and the solution to equation (2), the walker’s step-to-step dynamics can be studied as a hybrid Poincaré map [30], **F**, with state 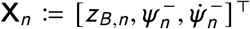 and input parameter vector 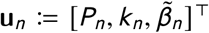 at step *n*:

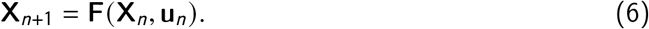

A period-1 gait (i.e., that repeats every step) corresponds to a fixed p oint 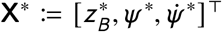 of the map **F** provided there is 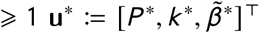 satisfying:

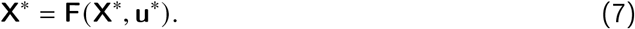

For a given periodic body state **X**^∗^ in the lab frame, there can be (infinitely) many input parameters **u**^∗^, and vice versa. Thus, every period-1 gait (**X**^∗^, **u**^∗^) is uniquely specified in t he six-dimensional (6D) *extended* state space consisting of body states and inputs.

### 2.2 Lateral body position and step width regulation

We impose a multiobjective task-level stepping regulator [6] that aims to achieve an *a priori* fixed body position 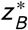 and step width *w* ^∗^ simultaneously by choosing optimal lateral foot placement of the walker (figure 2) at each walking step. At the task level, the regulated o bservables {*z*_*B*_, *w*} are related to the “end effector” observables, i.e., foot placements {*z*_*L*_, *z*_*R*_}, as:

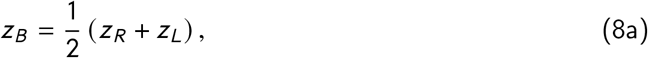

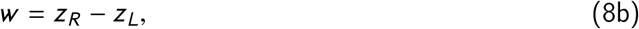

where *w* ⩾ 0, if there are no crossover steps.

The multiobjective regulation template selects the walker’s next lateral foot placement as a weighted combination of two predicted foot placements, one that optimally achieves the step width goal *w* = *w* ^∗^, and the other the body position goal 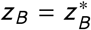 [6]. Let the walker’s right leg be its stance leg during step *n* (figure 2) so that

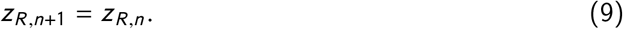

Then, at the (*n* + 1)^st^ heel strike, the regulator places the walker’s left foot laterally at 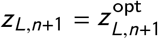, where

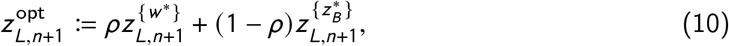

in which 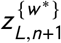 is the foot placement corresponding *w*_*n*+1_= *w* ^∗^ alone, 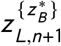 is the foot placement corresponding 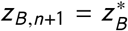 alone, and the mixture parameter *ρ* ∈ [0, 1] is kept fixed from step-to-step [6]: *ρ* = 1 corresponds to 100% *w* regulation and *ρ* = 0 to 100% *z*_*B*_ regulation. Using equations (8) and (9), we can write:

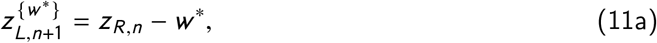

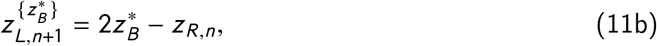

provided the walker can apply inputs that achieve the respective uniobjective targets *w* ^∗^ and 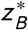 exactly. Substituting equations (11) in equation (10), we get the optimal left step as:

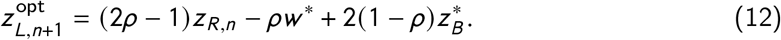

The resulting weighted foot placement 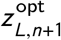 (equations 10 and 12), in fact, minimizes the following task-level cost exactly:

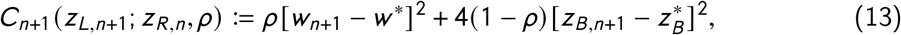

where *w*_*n*+1_ and *z*_*B,n*+1_ are functions of *z*_*L,n*+1_ (equation 8), and the optimal cost is (equations 11-13):

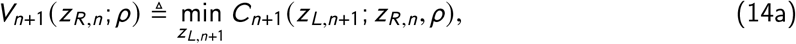

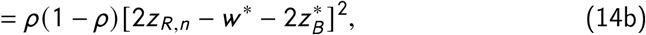

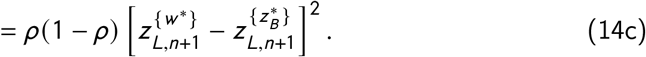

Similarly, for the left stance leg during step *n*, we obtain the walker’s right foot placement at 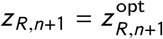, where

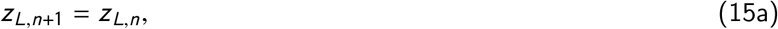

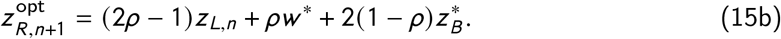

Using equation (8) together with equations (9), (12), and (15), yields linear fluctuation dynamics of the task-level observables {*z*_*B*_, *w*} under multiobjective regulation:

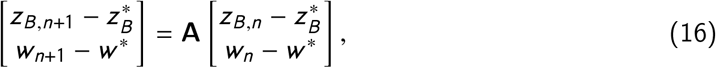

where

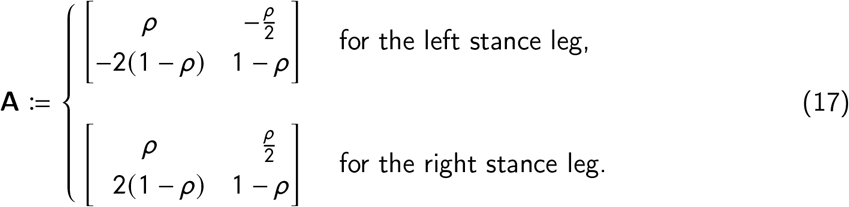

Since the matrix **A** only depends on *ρ*, linear task-level stepping dynamics (equation 16) is decoupled from body-level dynamics (equation 6) in the mechanical state space. However, the task-level observables [*z*_*B*_, *w*]^⊤^ (or, [*z*_*L*_, *z*_*R*_]^⊤^) are nonlinear functions of the mechanical state and, moreover, the dynamics in the walker’s state space itself is nonlinear.

Implicit in the successful realization of the task-level fluctuation dynamics (equation 16) is the requirement that the walker find valid inputs **u**_*n*_ for state **X**_*n*_ at each step (equation 6) so that the optimal foot placement can be achieved (equations 12 and 15b) or, equivalently, so that the global minimum of the multiobjective cost can be found (equations 13 and 14). We next show that not only is this possible, but also that the walker’s body-level dynamics is, in a certain sense, stable under this multiobjective regulation.

## 3 Do task-level regulators yield stable body-level dynamics?

We first find period-1 gaits (equation 7) of the lateral walker consistent with task-level regulation. The targeted fixed point, 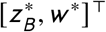, of the task-level dynamics (equation 16) directly determines two of the state variables, 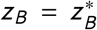 and 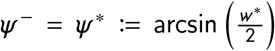 (figure 2), of any period-1 gait. Moreover, at the period-1 gaits, 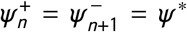 (equation 1a) and, since the walker’s stance phase is conservative (the energy 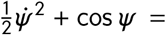 constant, via equation 2a), we have 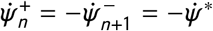, so that the push-off (equation 1b) satisfies:

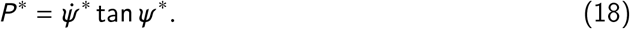

Thus, for a given 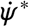 (or *P* ^∗^), one can obtain hip spring parameters 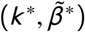 that form a curve Γ in the 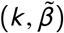 -plane representing the locus of period-1 gaits, where

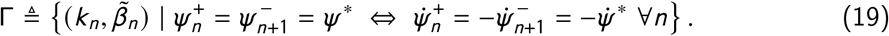

The set Γ is thus fully determined by 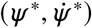 and is independent of the mixture parameter *ρ* (equation 10). Therefore, Γ yields the walker’s period-1 gaits irrespective of the task-level regulation (equation 16) or lack thereof (as for the unregulated, open-loop walker). This fact demonstrates that the decoupling between task and body level fluctuation dynamics, identified in the previous section, extends to period-1 fixed points themselves.

To further examine the set of period-1 gaits, we carried out numerical analysis for the multiobjective goal 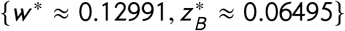 This yields a 1D manifold, ℳ_*x*_, of period-1 gaits (fixed points **X**^∗^, equation 7), in the state space (figure 3*a*), along a line segment 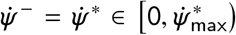 with 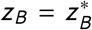, and *ψ* ^−^ = *ψ*^∗^ = 0.065, where the maximum angular rate (equations 4 and 5) is

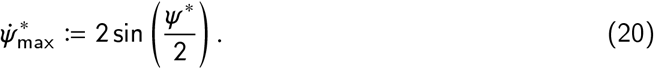

**Figure 3.**
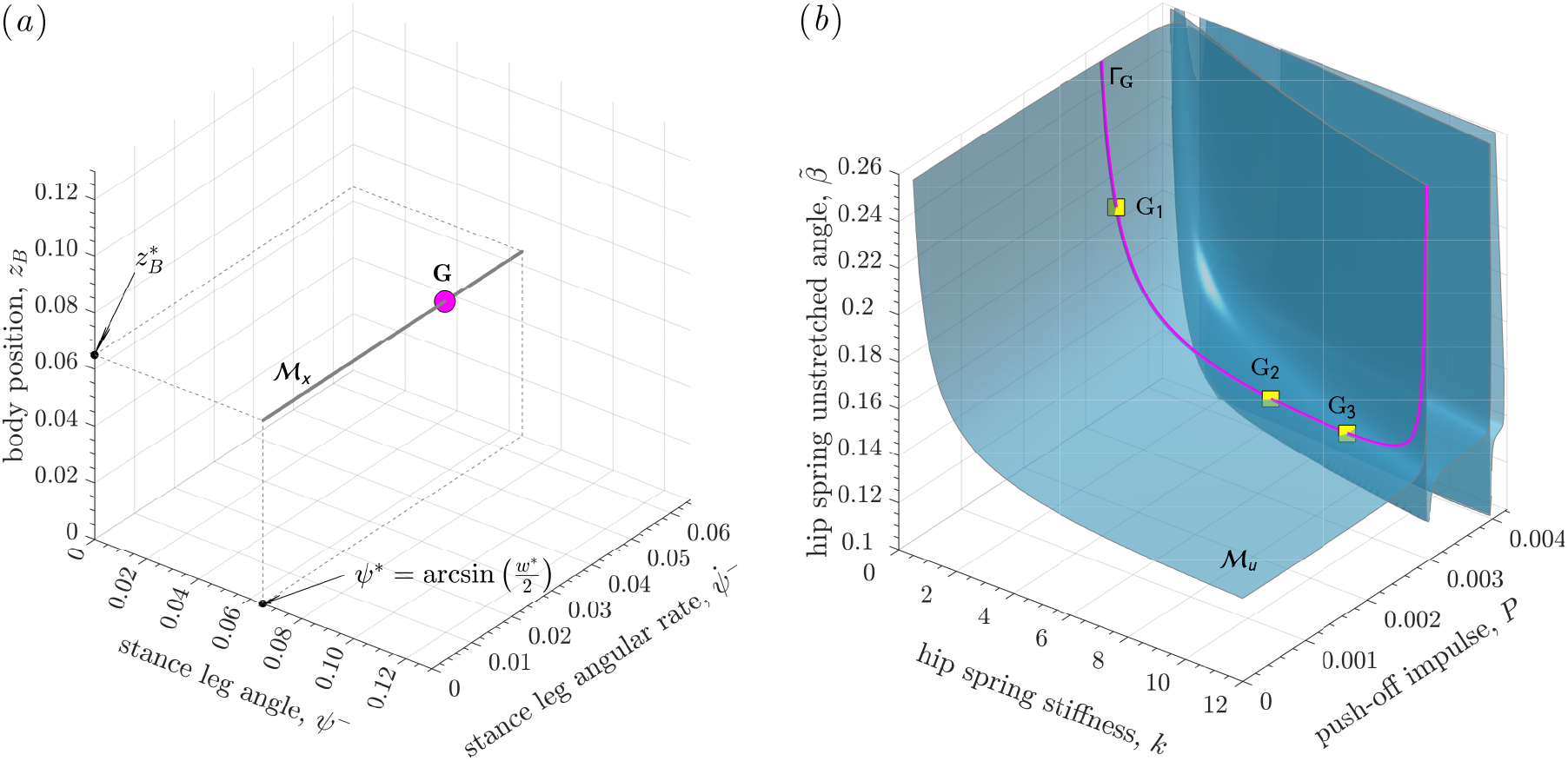
Period-1 gaits (equation 7) in the 6D extended state space comprised of (*a*) 3D mechanical state space and (*b*) 3D input space. For task-level multiobjective goal 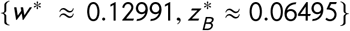, there is a 1D continuum of period-1 gaits with mechanical states on ℳ_*x*_ at 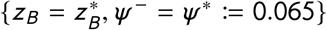 ((*a*), *thick straight line segment*). For every point on ℳ_*x*_ (e.g., point **G** at 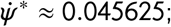 *circle*) there is ⩾ 1 curve (equation 19) of inputs in the 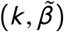 -plane (e.g, (*b*), Γ_**G**_ at 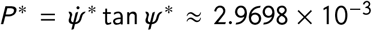; *thick curve*), each point on which yields a period-1 gait with same mechanical state. The points {G_1_, G_2_, G_3_} ∈ Γ_**G**_ (*squares*) represent three such gaits. The collection of all such curves forms the manifold ℳ_*u*_ (*b*). Thus, there is a 2D goal equivalent manifold 𝒢_P1_ (equation 26) in the 6D extended state space, every point in which is a period-1 gait achieving *w* = *w* ^∗^ and 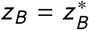.

Correspondingly, we get a range of push-offs 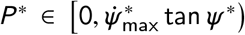 from equation (18). We estimated curves Γ (equation 19) for each 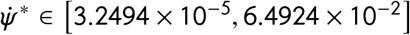, with increments of 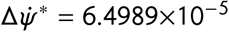, using a numerical continuation procedure. This yields the 2D manifold ℳ_*u*_ corresponding to period-1 gaits in the region 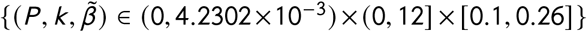 of the input space (figure 3*b*).

### 3.1 Goal equivalent manifold of period-1 gaits

A hallmark of biological motor performance is *equifinality* [2], which states that a given goal-directed movement can be executed in many, typically an uncountable infinity of, ways. While this makes its mathematical formulation ill-posed, it is an essential feature of biological movement that makes it adaptive and resilient in the face of physiological noise, external disturbances, changing environmental conditions, and injuries.

Goal-directed movement tasks can be defined using a *goal function*, **g**, of task-level observables (e.g., joint angles, or end effector positions and velocities) such that the zeros of **g** correspond to perfect task execution [19]. This set of task solutions defines a *goal equivalent manifold* (GEM), G, in the space of observables [31, 32]. In experimental applications, the goal function is generally unknown *a priori* but is a testable hypothesis on the movement [4–6, 33]. Here we demonstrate theoretically that the GEM has a dual representation in the system’s extended state space, which is the *combined* space of mechanical states *and* input parameters.

For lateral body position and step width regulation with goals 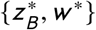, respectively, the goal function is expressed most simply as

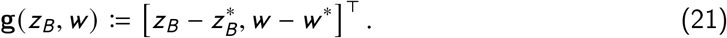

Thus, in the plane of task-level observables *z*_*B*_ and *w*, perfect execution of position and step width tasks are trivially specified by the lines 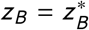 and *w* = *w* ^∗^, respectively, which define GEMs for position and step width goals *individually*. Using equations (8), these individual *z*_*B*_ and *w* GEMs get mapped to lines in the (*z*_*L*_, *z*_*R*_) stepping plane, used to display data from [6] in figure 1*c*. The *z*_*B*_ and *w* goals can be *simultaneously* achieved only at the point 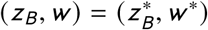 Here, we show how this “single point GEM” at the task level can be generated by the lateral walker.

We here assume that stepping regulation aims to achieve task-level goals 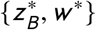 exactly at the next step. We thus obtain a “1-step” GEM, 𝒢_1_, defined as:

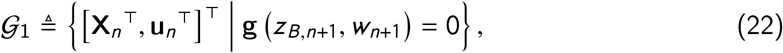

where the goal function **g** is from equation (21), but its arguments {*z*_*B,n*+1_, *w*_*n*+1_} are understood to be functions of the current extended state, (**X**_*n*_, **u**_*n*_). The 2 × 6 Jacobian matrix of **g**(*z*_*B,n*+1_, *w*_*n*+1_), which maps small deviations in states and inputs to deviations at the task level [19], is then

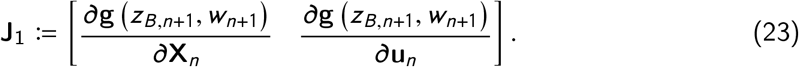

Generically, **J**_1_ has full rank (i.e., rank two) and, thus, the GEM 𝒢_1_ has co-dimension two, meaning that it is a 4D manifold in the 6D extended state space.

However, even when restricted to extended states (**X, u**) permitting a viable step [7], almost no points in 𝒢_1_ correspond to steady state gaits. Instead, under the dynamics of the system, extended states in 𝒢_1_ will evolve to whatever asymptotically stable gaits are permitted. Here, we focus on purposeful, steady walking, which we interpret as requiring that task-level goals be achieved via period-1 steady-state gaits. Thus, we define a “period-1” GEM, 𝒢_P1_, as the subset of all state-input pairs in the GEM 𝒢_1_ (equation 22) that also yield period-1 gaits (equations 6 and 7):

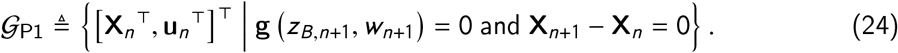

Provided such period-1 gaits exist, the corresponding 5 × 6 Jacobian matrix,

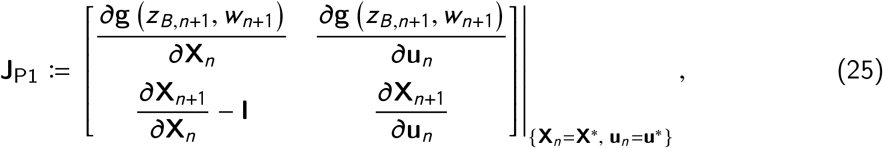

where **I** is the identity matrix, generically has rank four. Thus, the GEM 𝒢_P1_ has co-dimension four, so it is a 2D manifold in the extended state space. Equivalently, using the discussion surrounding equation (19), 𝒢_P1_ can be defined by

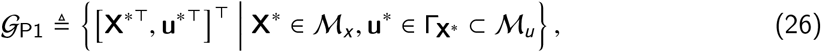

where Γ_**X**∗_ is the specific Γ curve of equation (19) (e.g., Γ_**G**_ in figure 3*b*) corresponding to the period-1 state **X**^∗^ (point **G** in figure 3*a*). Indeed, the sub-manifolds ℳ_*x*_ and ℳ_*u*_ (figure 3) are the projections of the GEM 𝒢_P1_ to the state and input space, respectively.

It is thus evident that there are uncountably many input parameters for any given period-1 gait that can equally achieve the task-level goal, as long as the walker remains viable [7]. That is, the lateral walker exhibits equifinality at the *body* level even though all such solutions are mapped into the point 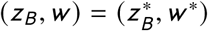 at the *task* level.

### 3.2 Implementing the multiobjective regulator

To implement optimal lateral foot placement (equations 12 and 15b) in code, we found it convenient to use the relationship of equation (8b) to optimize the step width:

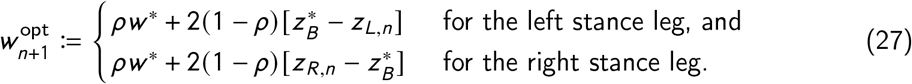

In each of these cases, at step *n* the walker’s regulator must select input parameters 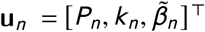 to pair with the current state **X**_*n*_ so that it can generate the next step. This is, again, mathematically ill-posed due to the existence of infinitely many solutions.

For our simulations, we set **u**_*n*_ to 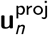, defined as the orthogonal projection of the walker’s previously applied input, **u**_*n*−1_, onto the 2D target optimal-step-width manifold ℳ_*u,n*+1_ in the input space corresponding to state **X**_*n*_ :

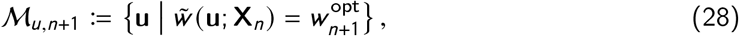

where 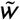 is the step width considered as a function of the inputs with the current state **X**_*n*_ taken as a known parameter. In short, ℳ_*u,n*+1_ is the set of all inputs guaranteeing that 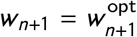 (equations 27) at the next step; **u**_*n*_ ∈ ℳ_*u,n*+1_ is a closest such input to **u**_*n*−1_. Note that, in general, ℳ_*u,n*+1_ < ℳ_*u*_ (figure 3*b*) because 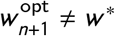. Given **X**_*n*_, 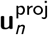was iteratively estimated until the magnitude of the smallest angle between the vector 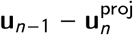 and the normal to ℳ_*u,n*+1_ at 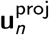 was less than 0.1°. This geometrical construction implies that our regulator implementation results in a step-to-step map acting on the space of state-input pairs, (**X, u**).

### 3.3 Semistability of the walker under multiobjective regulation

Task-level regulation does not in itself guarantee the existence of stable gaits. Thus, we first simulated the walker’s trajectories with *ρ* = 0.9 (90% step width and 10% body position regulation; equations 10 and 13), as estimated in humans (figure 1;[6]). Typical trajectories of all six variables in the system’s extended state space, starting from different initial states near a given period-1 gait, are shown in figure 4, in which two time scales are visible. In figure 4 (*left column, top two*), task-level regulation rapidly (in under 50 steps) drives *z*_*B*_ and *ψ* ^−^ very close to their target values, thus placing the mechanical state, **X**, nearly on ℳ_*x*_ (figure 3*a*) and its extended state, (**X, u**), very close to the *targeted subset* of GEM 𝒢_1_ (equation 22), i.e., the portion of 𝒢_1_ on which 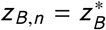and 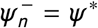and, hence, the task-level goal function (equation 21) is **g** ≈ 0 for all subsequent steps.

**Figure 4.**
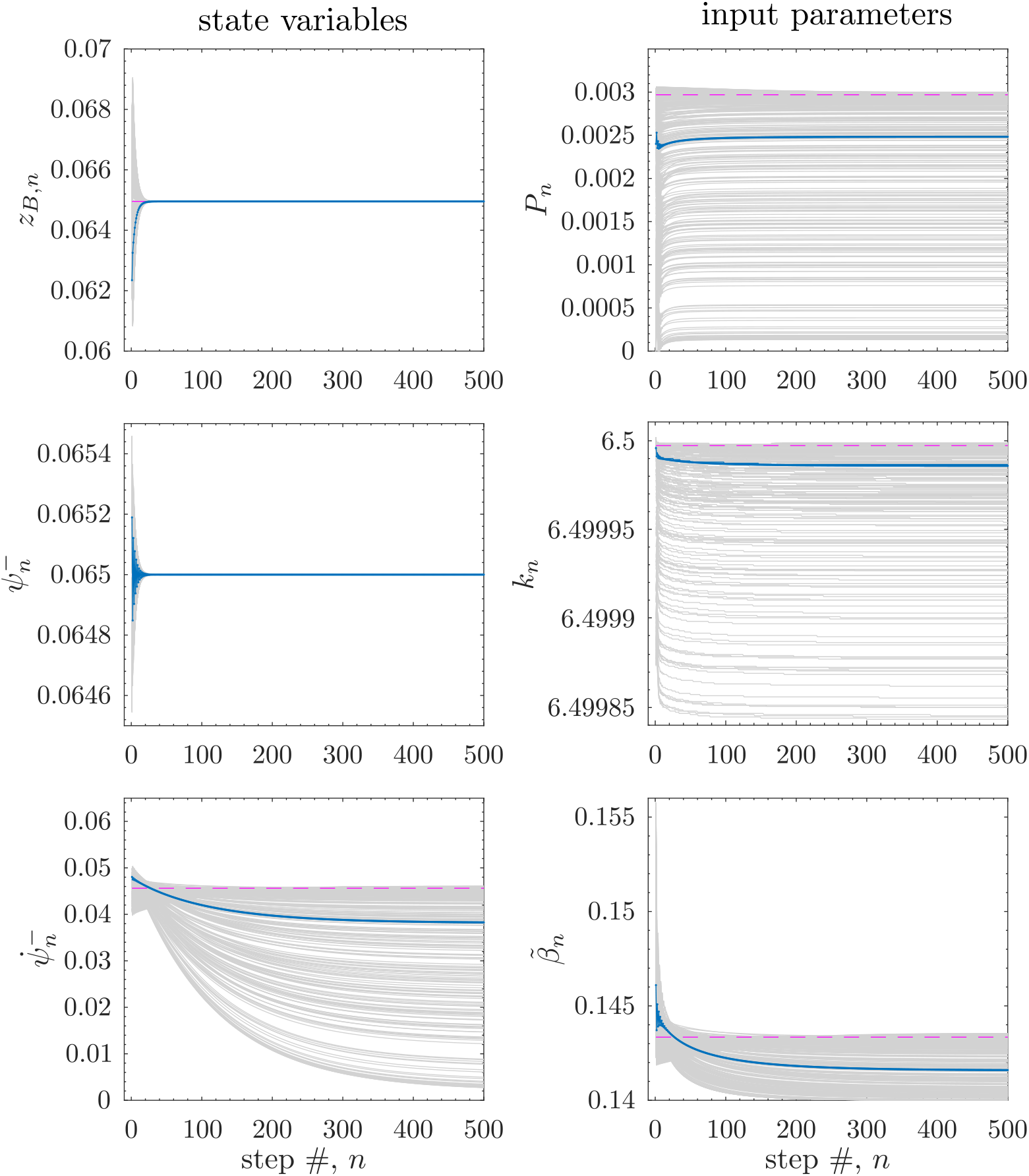
Semistable dynamics of period-1 gaits in the regulated system with *ρ* = 0.9: ensemble of time series (*thin gray curves*) of state, **X** (*left column*), and input, **u** (*right column*), starting from 475 distinct initial mechanical states **X**_1_ sampled uniformly randomly from surface of an ellipsoid around state **X** = **G** (figure 3*a*):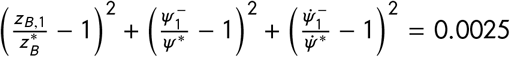. All trajectories used input **u**_0_ = 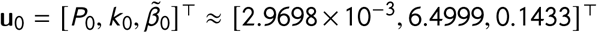 (*right column, broken horizontal lines*), point G_2_ in figure 3*b*. Representative time series (*highlighted thick blue curve*) typifies semistability of regulated period-1 gaits: trajectories asymptotically approach distinct, Lyapunov stable period-1 gaits with mechanical states lying on ℳ_*x*_ (figure 3*a*). Directly regulated state variables (*left column, top* and *middle*) rapidly approach their target values; the unregulated state variable (*left column, bottom*) more slowly approaches a point on ℳ_*x*_, typically different from the reference point **G**. At the same time, input trajectories (*right column*) approach distinct values on ℳ_*u*_ (figure 3*b*), typically different from G_2_.

However, at this point, the input **u** is not yet near ℳ_*u*_ (figure 3*b*) and the system has yet to reach steady state. Further transients, particulary visible in the unregulated variables (figure 4 *left column, bottom* and *right column*), occur until trajectories settle (at about 500 steps) on period-1 gaits in GEM 𝒢_P1_ (equation 26), with different 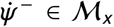 and distinct steady state inputs in ℳ_*u*_ (figure 3*b*). The latter stage of transients is an example of *self motion*, body-level dynamics that do not affect task-level performance [34, 35], consisting in this case of trajectories along the targeted subset of 𝒢_1_ that asymptotically approach the 2D 𝒢_P1_ while simultaneously satisfying task goals.

These numerical results demonstrate that at least a subset of 𝒢_P1_, the GEM containing period-1 gaits, is *semistable* [24–27]: trajectories starting from perturbations off of any gait in this set remain bounded and asymptotically return to other gaits embedded within it. A direct consequence of equifinality and the non-uniqueness of steady states, semistability is less constraining than the asymptotic stability of limit cycle walking. At least for disturbances in some neighborhood of 𝒢_P1_, the walker is guaranteed to return to a period-1 gait that, while typically different than the original, lies in the same GEM and so will achieve the same task goals. Indeed, using equation (14), one can show

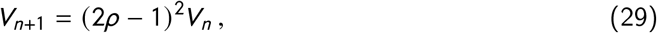

so the minimized cost *V*_*n*_ evaluated along trajectories (as, e.g., in figure 4) decreases monotonically for 0 < *ρ* < 1 (figure 5). Thus, *V*_*n*_, defined in terms of task-level observables that themselves are functions of the walker’s state, behaves like a *Lyapunov function* [24–27] with respect to the targeted subset of 𝒢_1_ (equation 22). This is consistent with the semistability of the walker’s period-1 gaits on the attracting submanifold of 𝒢_P1_ ⊂ 𝒢_1_ (equation 26).

**Figure 5.**
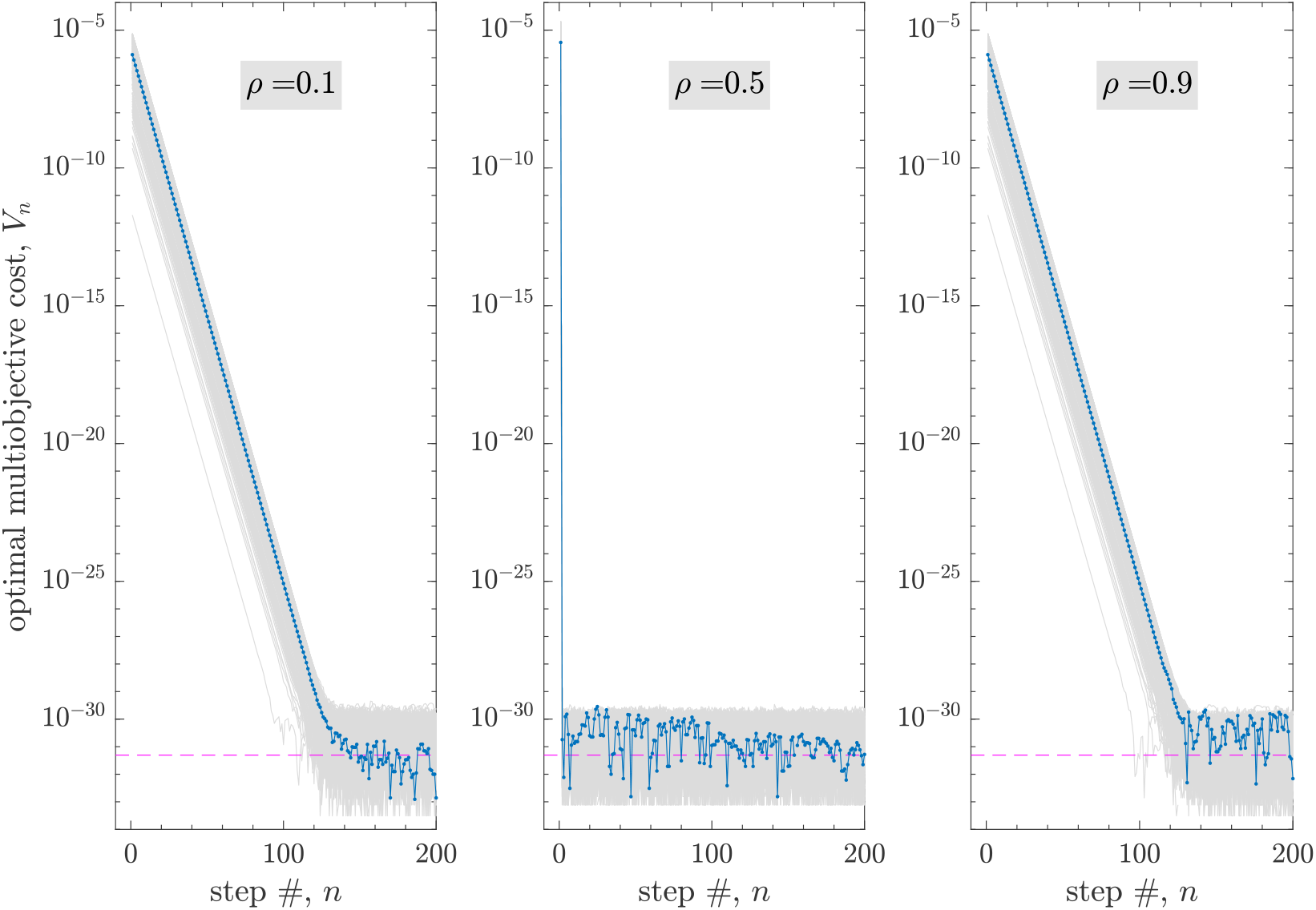
Monotonic decrease of the optimal multiobjective cost *V*_*n*_ (equation 14) along the task-regulated walker’s deterministic trajectories initialized near gait G_2_ (figure 3). For *ρ* = 0.9 stepping trajectories shown in figure 4. For *ρ* * 0.5, *V*_*n*_ hits a “noise floor” near *ε*^2^ (*broken horizontal line*) after exponential decay (*ε* ≈ 2.204 × 10^−16^, eps in MATLAB), with semilog slope 2 log |2*ρ* − 1| (from equation 29). This indicates an asymptotic approach to the targeted subset of GEM 𝒢_1_ (equation 22) that holds 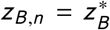and 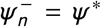(compare with figure 4, *left column, top two*). For *ρ* = 0.5, *V*_*n*_ = 0 (within numerical precision) in two steps, indicating finite-time convergence. Irrespective of *ρ*, trajectories asymptotically approach the semistable subset of the GEM 𝒢_P1_ of period-1 gaits (equation 26).

### 3.4 Lateral walker with motor noise: fluctuation dynamics

To compare the integrated template’s lateral stepping dynamics with human data, we simulated the system with “motor” noise, modeled as independent Gaussian noise of amplitude *σ*_*m*_ on the input at step *n*:

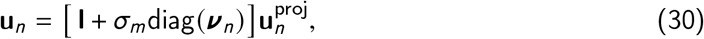

where 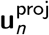(equation 28 and surrounding discussion) is the deterministically optimal input, **I** is the identity matrix, ***ν*** _*n*_ is a vector of standard normal random variables, and diag(***ν*** _*n*_) is a diagonal matrix with principal diagonal ***ν*** _*n*_.

With the mixture parameter again set to *ρ* = 0.9, the stochastically perturbed walker with 1% Gaussian motor noise (*σ*_*m*_ = 0.01) was able to walk (i.e., to remain viable [7] for) at least 500 steps. Ensembles of time series obtained from 475 stochastic simulations, all starting from the same period-1 gait, are shown in figure 6. Deviations in the regulated variables, *ψ* ^−^ and *z*_*B*_, both display significant diffusion, but relatively low drift: instead, they oscillate closely around their target values (recall that *w* = 2 sin *ψ* ^−^). Fluctuations in both variables appear to settle into a stochastic steady state after roughly 200 steps. However, steady-state *ψ* ^−^ fluctuations have visibly smaller temporal correlations (i.e., their random oscillations are much more rapid) than those of *z*_*B*_ : as discussed in more detail below, this arises from the much stronger weight (*ρ* = 0.9) given to *w* regulation in these simulations.

**Figure 6.**
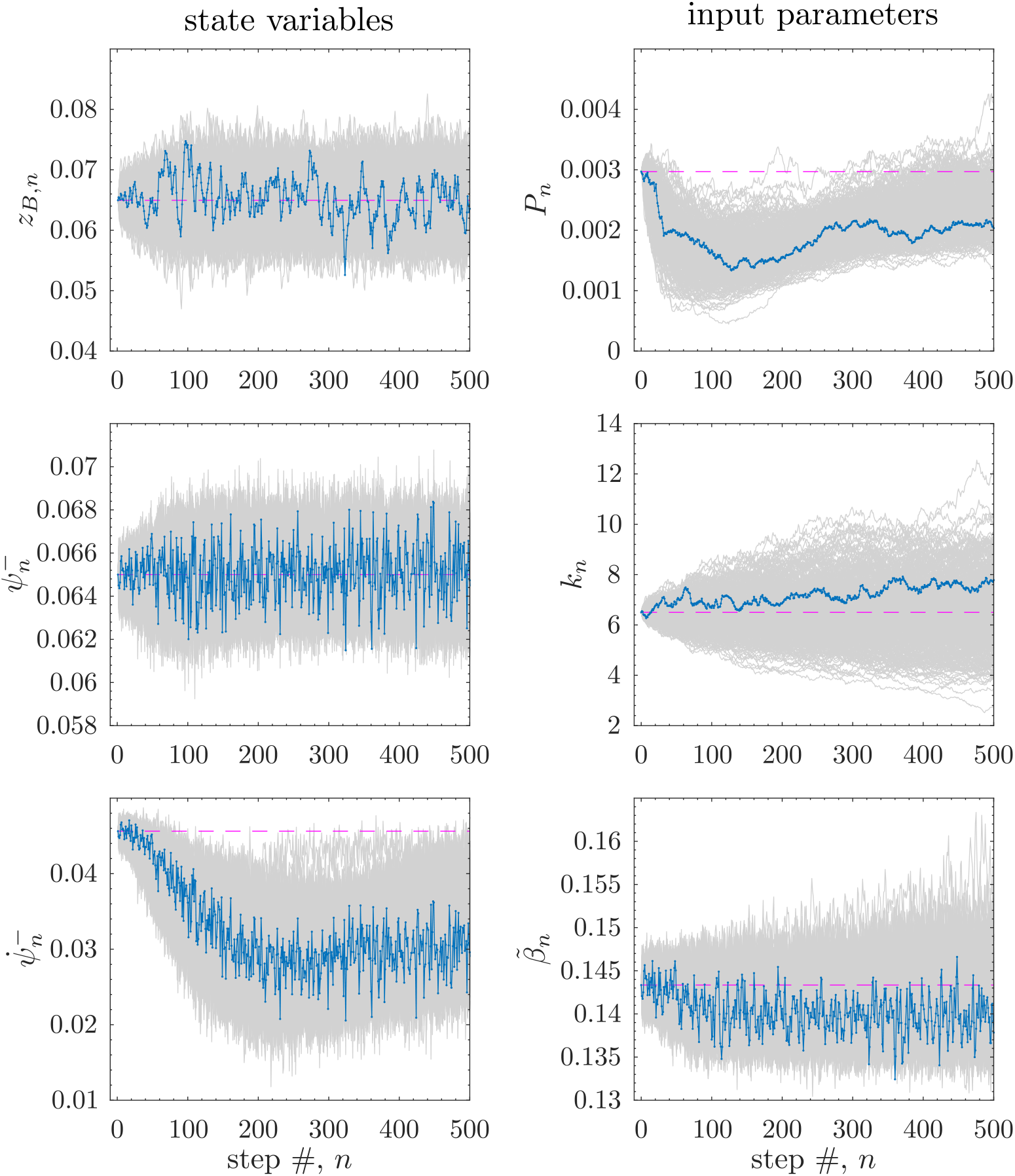
Lateral walker with simulated motor noise: ensembles (*thin gray curves*) of state (*left column*) and input (*right column*) time series with *σ*_*m*_ = 1% Gaussian noise (equation 30) added to optimal inputs at each step, with *ρ* = 0.9. Initial conditions (*broken horizontal lines*) corresponded to gait G_2_ (figure 3); 475 stochastic realizations (time series) were generated using distinct seeds to a pseudorandom number generator. A representative time series (*highlighted thick blue curve*) demonstrates typical behavior under stochastic perturbations: mechanical state trajectories are regulated transverse to the GEM and so oscillate close to ℳ_*x*_ (figure 3*a*) after a diffusive transient (*left column, top* and *middle*); however, there is drift away from the initial period-1 orbit along ℳ_*x*_ (*left column, bottom*) and in the input space (*right column*; also see ℳ_*u*_, figure 3*b*). The set of period-1 gaits on the GEM on which trajectories settle (say, at step 500) with high probability depends on initial conditions. Consistent with 90% *w* regulation (*ρ* = 0.9), *ψ* ^−^ time series (given *w* = 2 sin *ψ* ^−^) shows significantly less persistence of fluctuations compared to *z*_*B*_ (also see figure 7).

In contrast, the time series ensembles for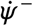, *P*, *k*, and 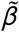, which are not directly regulated, display both diffusion and drift. The drift is an example of *stochastic* self motion along 𝒢_1_, during which, in contrast, the regulated variables fluctuate with approximately zero mean about their targeted values. Thus, the mechanical state drifts primarily in 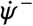, i.e., along the manifold ℳ_*x*_ (figure 3*a*), consistent with the semistability of the deterministic walker (figure 4), as it settles on a stochastic steady state near a new period-1 gait. The drifting behavior is a stochastic transient that lasts for hundreds of steps, longer than required to achieve approximate steady state at the task level (variables *ψ* ^−^ and *z*_*B*_). One finds that, again consistent with semistability, the steady-state probability distributions of these drifting variables depend on their initialization in the extended state space (figure 3; equation 26).

The temporal correlation differences observed in the plots of *ψ* ^−^ and *z*_*B*_ (figure 6) can be quantified using detrended fluctuation analysis (DFA; [36–38]), which yields the scaling exponent *α* : *α* > 0.5 indicates statistical *persistence*, meaning that a fluctuation at one step is likely to be followed by a fluctuation of the same sign at the next step, whereas 0 < *α* < 0.5 indicates *anti* persistence, so fluctuations at sequential steps are likely to alternate in sign; *α* = 0.5 for uncorrelated noise and *α* ⩾ 1.5 for unbounded random walks.

In experiments with human participants [4, 5, 33, 39–43], the DFA exponent provides a signature of motor regulation: tightly regulated task-level variables tend to have *α* ≈ 0.5, whereas unregulated variables have 0.5 ≪ *α* < 1.5. This “tightness-of-control interpretation” [43] of DFA suggests an hypothesis on the organization of the human motor regulation system that is, however, difficult (if not impossible) to study only through experiments. Even the most clever of experimental protocols have limited access to internal parameters of the neuromotor and biomechanical systems responsible for movement and, even when there is such access, the relevant parameters cannot typically be adjusted.

We thus estimated *α* for noisy time series simulated with different values of the mixture parameter *ρ* to see if its effect on the fluctuation properties of the observables {*w*, *z*_*B*_, *z*_*L*_, *z*_*R*_} conformed with the tightness-of-control hypothesis. Recall these observables are not independent (equations 8) and the regulator explicitly minimizes costs in *w* and *z*_*B*_, weighted by *ρ* (equation 13). For *ρ* = 0.9 (figure 7 *right*), the strongly regulated variable, *w*, has *α* [*w*] ≈ 0.5, whereas the weakly regulated *z*_*B*_ has much higher values (*α* [*z*_*B*_] ≈ 1). Furthermore, the lower persistence of *w* relative to *z*_*B*_ (*α* [*w*] ≪ *α* [*z*_*B*_]) quantifies the apparent lower temporal correlation of *w* with respect to *z*_*B*_, visible in the time series of figure 6 (*left column, top two*). In contrast, the individual step positions, *z*_*L*_ and *z*_*R*_, which are not directly regulated, have high persistence, with DFA exponents comparable to *α* [*z*_*B*_]. Taken together, these results closely mirror those found in experiments with walking humans (figure 1*d*).

**Figure 7.**
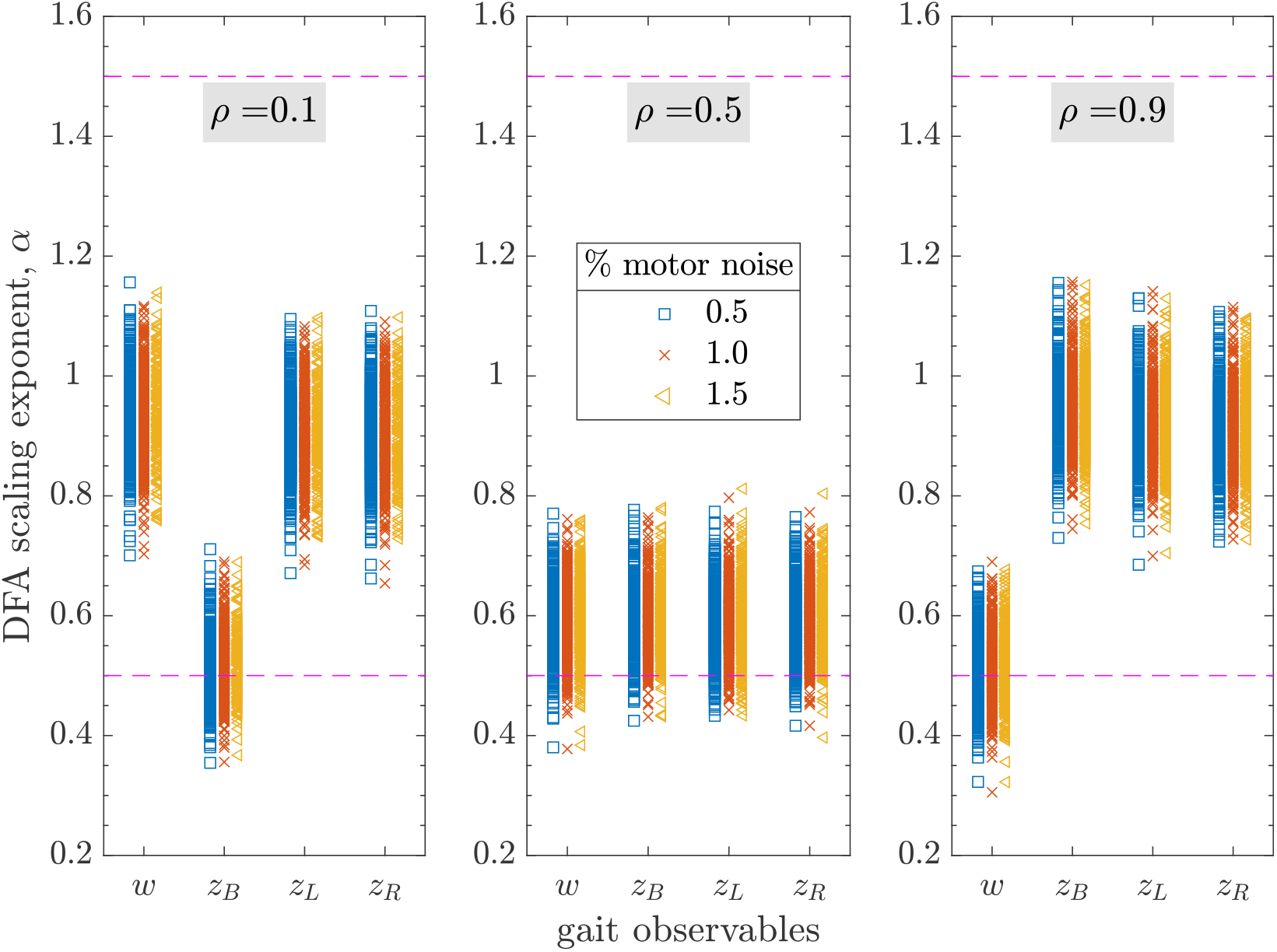
Detrended Fluctuation Analysis (DFA) results for the regulated walker with simulated motor noise and mixture parameter *ρ* ∈ {0.1, 0.5, 0.9}. We computed scaling exponents *α* (*markers*) for noisy time series (475 trials; 500 steps per trial) of stepping observables {*w*, *z*_*B*_, *z*_*L*_, *z*_*R*_} at three motor noise levels *σ*_*m*_ ∈ {0.5, 1, 1.5}%: for reference, *α* = 0.5 for white noise and *α* = 1.5 for a Brownian random walk (*broken lines*). All time series were initialized at gait G_2_ (figure 3), as in figure 6 for *ρ* = 0.9. Consistent with the “tightness-of-control interpretation” of *α*, fluctuations in step width become more persistent (*α* [*w*] increases) and those in the body position become less persistent (*α* [*z*_*B*_] decreases) as the degree of step width regulation *ρ* decreases from 0.9 to 0.1. Fluctuations in all observables show uniformly low persistence when both *w* and *z*_*B*_ are regulated to the same degree (*ρ* = 0.5). Humans walking on a treadmill choose their lateral foot placements in a way that resembles multiobjective regulation with *ρ* ≈ 0.9 ([6]; figure 1*d*). Note that DFA *α* is not significantly affected by noise strength *σ*_*m*_.

Figure 7 (*left*) shows the corresponding DFA results for *ρ* = 0.1. Here, the relative degree of persistence swaps between the two variables: for *z*_*B*_, which is strongly regulated, *α* [*z*_*B*_] ≈ 0.5, whereas *α* [*w*] ≈ 1. Finally, when both variables are equally regulated (*ρ* = 0.5, figure 7 *center*), the persistence of fluctuations is comparably low for all the variables. These simulation results support the tightness-of-control hypothesis: as the regulation shifts from predominantly weighting *w* to *z*_*B*_, the DFA *α* for both undergoes a corresponding shift, as predicted. The uniformity of *α* values across all variables when *ρ* = 0.5 is a new observation, but not completely unexpected, as discussed below.

Typical scatter plots of stepping data from the stochastically perturbed walker are shown in figure 8, in which columns correspond to simulations with *ρ* ∈ {0.1, 0.5, 0.9}. The top two rows in each column present “direct control” analyses ([6]; figure 1*a,b*) of the simulated data: for perfect regulation, meaning that a deviation from the target value is, on average, eliminated at the next step, the regression line in such plots would have slope −1. In the right-most column of figure 8, for which *ρ* = 0.9, we see that the direct control plot for *w* indeed has a regression slope ≈ −1, whereas the slope for the *z*_*B*_ is near zero. This is entirely as expected, given that in this case *w* and *z*_*B*_ are strongly and weakly regulated, respectively. Importantly, these results are consistent with the DFA results of figure 7 (*right*) for the same value of the mixture parameter. Furthermore these simulation results again mirror those obtained in experiments with human participants (figure 1*a,b*).

**Figure 8.**
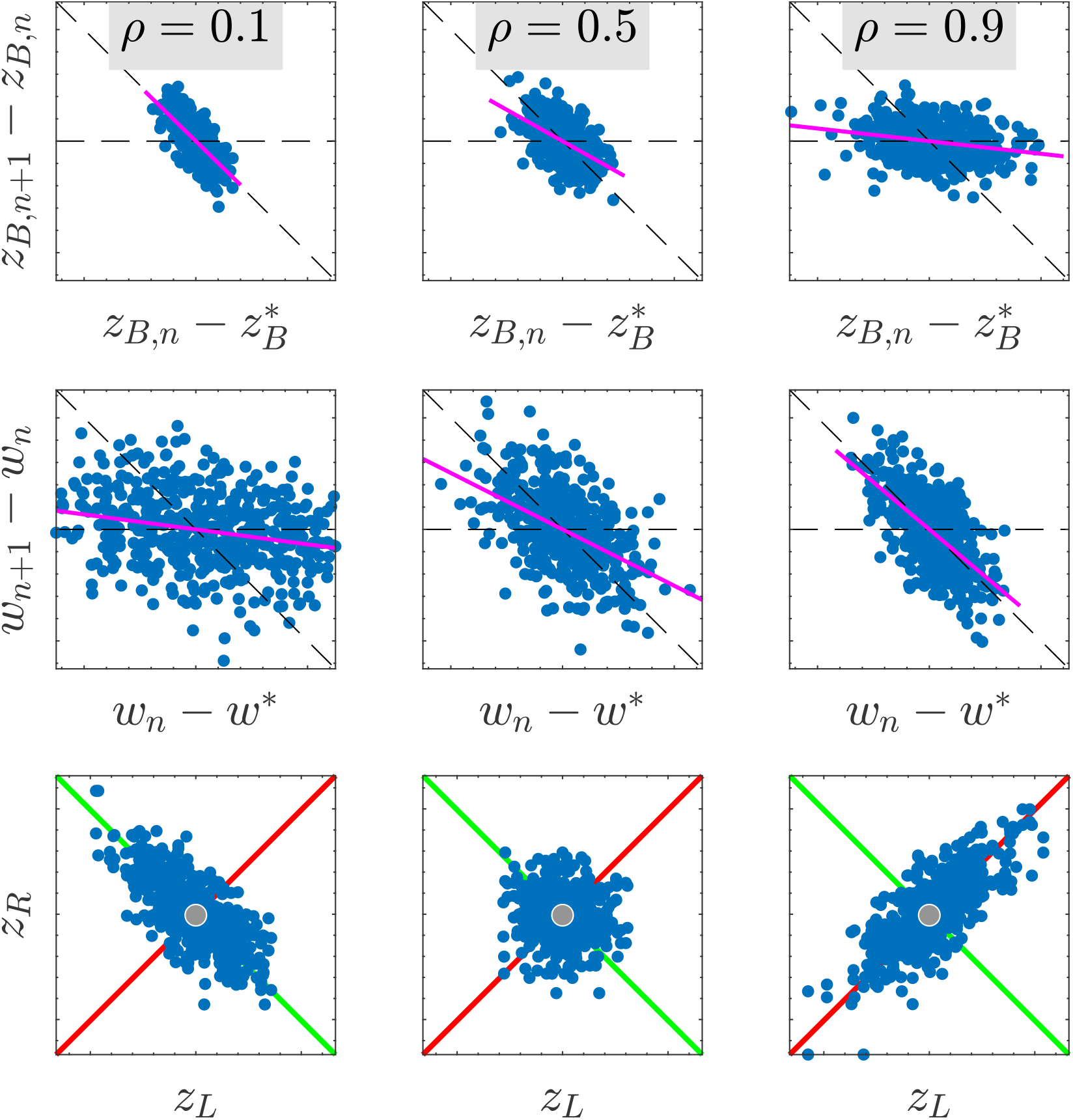
Typical step-to-step fluctuation data (*dots*) of lateral body position (*z*_*B*_), step width (*w*), and foot placements (*z*_*L*_, *z*_*R*_) of the stochastically perturbed (1% motor noise) regulated walker for *ρ* ∈ {0.1, 0.5, 0.9}. Compare each *column* (at a given *ρ*) to a typical experimental trial in human walking (figure 1*a*-*c*). As the degree of step width regulation *ρ* decreases, the persistence of step-to-step fluctuations in *z*_*B*_ also decreases (*regression line* slope changes from ≈ 0 to ≈ −1; *top row*), while that in *w* increases (*regression line* slope changes from ≈ −1 to ≈ 0, *middle row*). Same walking trial plotted in the stepping plane (*bottom row*) shows increase in fluctuations in *z*_*B*_ and simultaneous decrease in fluctuations in *w* as *ρ* increases: *data ellipse* is oriented along 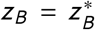 (*green line*) for *ρ* < 0.5 and along 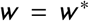 (*red line*) for *ρ* > 0.5. At *ρ* = 0.5, the variability structure of foot placements is isotropic. The stepping fluctuation dynamics and the variability structure for *ρ* = 0.9 (*right column*) resembles that observed in human data (figure 1*a*-*c*).

Varying *ρ* in our simulations gives results that similarly cohere with the DFA analyses of figure 7: in figure 8 (*top row*), as *ρ* varies from 0.1 to 0.9, the regression slope for the *z*_*B*_ direct control plots changes from ≈ −1 to ≈ 0, while the slope for the *w* plots (figure 8 *middle row*) exhibits the opposite change. Thus, DFA and direct control analyses provide complementary and consistent characterizations of regulation strength.

The bottom row of figure 8 displays stepping data in the (*z*_*L*_, *z*_*R*_) plane. As in the experimental plot of figure 1*c*, lines are included indicating task-level GEMs for 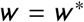 and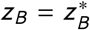, individually; the combined “GEM” for the multiobjective task is the point at their intersection. In all cases, we observe that principal axes of the data ellipses are approximately aligned with the single-variable lines, and the sample mean is near the intersection point. For *ρ* = 0.9, the major axis of the ellipse lies along the *w* GEM, as expected: strong *w* regulation minimizes the distance of steps from this line, whereas weak *z*_*B*_ regulation permits steps to spread along it. When *ρ* = 0.1, the strength of regulation reverses, and stepping data aligns with the *z*_*B*_ GEM.

When *w* and *z*_*B*_ are regulated to the same degree, at *ρ* = 0.5, the data ellipse (figure 8 *bottom row, center*) is close to a circle. This phenomenon has been observed in experiments in which people were asked to change lanes while walking [44, 45]: during steady walking before and after the lane change, stepping data conformed with the *w* -dominated *ρ* ≈ 0.9 case of figure 8 (*bottom row, right*; and as in figure 1*c*), whereas during the transition step, scatter plots were closer to circles, suggesting that participants regulated *w* and *z*_*B*_ nearly equally. This isotropy of the simulated stepping plane data when *ρ* = 0.5 provides a geometrical counterpoint to the DFA of the same gait observables for the same value of *ρ* (figure 7, *center*).

We conclude this section with a caveat. Here, we can interpret DFA from first principles: we know what variables are being regulated as parameters are varied, and so can show how this regulation correlates with estimates of *α*. However, care must be taken when using DFA exponents to make inferences about motor regulation strength from experimental data because completely passive, open loop aspects of a system impact DFA. It can easily be shown that *α* is functionally related to the local stability properties of a given steady state [31], whether or not the system is feedback controlled. Thus, while the tightness-of-control hypothesis says that if a particular variable is tightly regulated, then *α* ≈ 0.5, the converse need not be true. For this reason, tightness-of-control interpretations of experimental DFA results need to be grounded in the analysis of data from task-level variables with a plausible need for motor regulation and should, whenever possible, be checked against results for other variables and under conditions that might change such regulation. Finally, the resulting assessments should be checked against other empirical measures of correlation as well.

## 4 Discussion and conclusions

We introduced a simple lateral dynamic walker, with swing leg dynamics and three adjustable input parameters, to study how motor regulation affects frontal plane stepping (figure 2). The walker’s adjustable push off provides between-step actuation; its adjustable hip spring tunes within-step swing dynamics. Motivated by experimental observations and phenomenological models of human stepping regulation (figure 1; [6]), we imposed task-level multiobjective regulation targeting the walker’s optimal lateral foot placement at each step. The regulator prioritizes achieving a step width, vis-à-vis the walker’s lateral body position, to varying degrees by choosing a mixture parameter *ρ*. Our model thus integrated a lateral *mechanical template* [17], which captures fundamental mechanics of frontal-plane walking, with a lateral *motor regulation template* [6], an empirically verified model of how humans manipulate lateral foot placements in a goal-directed manner.

Our system provides a physically grounded example of the hierarchical control schema hypothesized in experiments [4–6, 19, 31–33, 44], in which an inner “control” loop responsible for movement execution serves as the plant for an outer “regulation” loop used to accurately achieve movement goals. In engineering, such cascaded architectures typify applications of *successive loop closure* (SLC) [9, 10], a design approach used in automation and robotics including, most relevantly here, autonomous vehicles [46–48]. SLC provides a useful framework for conceptualizing human movement on multiple grounds. First, the neurophysiology of the motor system possesses a similar hierarchical structure [11, 12], with an “inner” subsystem responsible for movement execution and an “outer” subsystem responsible for motor planning and coordination (see, e.g., [11, figure 4]). Second, SLC lends itself to the incremental development of cascaded models that, in principle, span all levels from “neurons to whole-body motion” [17]. A given integrated template might be “anchored” by further physiological and/or biomechanical inner loops: e.g., the hip spring in the template studied here might be “unpacked” into an inner loop involving reflex-mediated muscle groups [49]. Likewise, outer loops might be added representing perceptual/cognitive processes needed, e.g., for navigation and obstacle avoidance [50]. Finally, SLC conceptually parallels biological motor skill acquisition in which new motor capabilities are developed by exploiting previously learned, more basic abilities (e.g., children first learn to crawl, then stand, then walk, then run [51]).

Experiments with human participants [6] tested this general schema using hypotheses on error-correcting regulation at the task level, meaning, for walking, with respect to goals specified solely by stepping observables. This paper provides the first demonstration that such task-level regulation can arise from a physics-based dynamic walking system and, in turn, resolves two outstanding theoretical challenges. Arguably the most fundamental of these challenges concerns how empirical multiobjective linear models, in stepping observables alone, can so effectively capture observed stepping dynamics (e.g., [6]; figure 1). This may seem surprising given that, not only is the underlying biomechanical system undoubtedly nonlinear, but such models are estimated without reference to the walker’s mechanical state, thereby seeming to “ignore Newtonian dynamics”. However, our integrated template model precisely achieves this condition: the multiobjective regulator design yields linear step-to-step dynamics at the task level that are decoupled from the walker’s body-level nonlinear dynamics (see equations 16, 17, and 19 and surrounding discussion). Motor variability and regulation in the presence of equifinality [2] has only been studied experimentally using goal equivalent manifolds (GEMs) [19, 31] defined in the space of task-level observables [4, 6, 33]. Thus, a second challenge concerns how goal equivalence at the task level, which is directly observable in experiments, is reflected at the body level, which is not. Here, we demonstrated that the body-level GEM is a manifold in the system’s *extended state space* (the combined space of mechanical states and inputs). Our system has 3D mechanical state and input parameter vectors (section 22.1), giving a 6D extended state space. For an optimal stepping regulator specifying only target values of step width and lateral position, we obtained a 4D GEM, 𝒢_1_ (equation 22) in the extended state space. However, most points in 𝒢_1_ are not steady state gaits: we showed that the dynamics of the regulated system drives extended states onto 𝒢_P1_ (equation 26), a 2D submanifold of 𝒢_1_, that contains a continuum of period-1 gaits all satisfying stepping goals and, hence, is also a GEM.

We studied the dynamics in the extended state space by simulating the walker’s trajectories with *ρ* = 0.9 (90% step width and 10% body position regulation), consistent with models of lateral stepping regulation in humans (figure 1; [6]). We found that at least a subset of the GEM containing the walker’s period-1 gaits is *semistable* [24–27]. That is, if the system is perturbed away from any of the continuum of goal equivalent period-1 gaits in 𝒢_P1_, the transient behavior attracts the state back to 𝒢_P1_, but typically to a *different* period-1 gait. Thus, while the *individual* solutions are not asymptotically stable, at least a portion of the GEM *is*. Indeed, the walker’s minimized multiobjective cost at each step (equation 14a) behaves like a Lyapunov function (equation 29; [27]), demonstrating the asymptotic stability of the targeted subset of 𝒢_1_ on which the task goals are satisfied for all steps, and which itself contains 𝒢_P1_. This suggests semistability can be expected to be a property of systems with “GEM aware” optimal regulators of the type studied here.

The dynamics of the walker with simulated motor noise showed that it can remain viable [7] for several hundred steps, displaying stepping fluctuation statistics closely mirroring those observed in humans. Results were consistent with the “tightness of control” interpretation of the DFA exponent, *α* [43]: fluctuations in step width become more persistent, as quantified by *α*, and those in body position become less persistent as *ρ* decreases from 0.9 to 0.1. The degree of persistence in the task-level observables at *ρ* = 0.9 (figure 7) matches well with that observed in human experiments (figure 1*d*). Moreover, the walker’s step-to-step fluctuation dynamics and task-level variability structure for *ρ* = 0.9 (figure 8, *right column*) resembles that observed in human data (figure 1*a*-*c*). Step-to-step foot placements show least persistence and a nearly isotropic variability structure when *ρ* = 0.5 (i.e., with the same degree of step width and body position regulation), a phenomenon observed as people take transition steps during lane change maneuvers [44, 45]. Finally, even though the step-to-step task-level dynamics are strongly regulated, the walker’s stochastic trajectories at the body level show significant drift and diffusion of the mechanical states and inputs, consistent with the minimum intervention principle [52, 53] and the phenomenon of self motion [34, 35].

We found that semistable walking relative to a goal equivalent manifold of gait solutions was a natural consequence of task-level optimal regulation. Semistable walking is a less constraining paradigm than that provided by asymptotically stable limit cycles and, therefore, is more suitable for biological walking, which is characterized by its flexibility and adaptability. Moreover, semistability is consistent with the idea that humans can, at least in principle, remain viable (i.e., avoid falls) by rapidly switching between regulation strategies and corresponding goal equivalent sets of steady-state gaits [7]. Supported by our results, and the knowledge that equifinality is commonplace in human movement, we propose that optimal task-level regulation provides a conceptualization of walking as arising from an already-learned collection of viable stepping solutions between which humans can switch at each step by, e.g., changing task goals and/or cost functions. This allows for highly adaptive gaits that enable walkers to quickly and effectively respond to changing task requirements, environmental conditions, and external disturbances.

The regulated compass walker developed here provides a simplified model of lateral stepping observed in human gait. However, in the spirit of [17], referring to our model as a “template” emphasizes that it not only provides a simplified description of the system, but also serves as a “guide” or “target” for the control used by the entire, much more complicated organism into which the simplified model should be seen as being “anchored”. Our model’s features are biomechanically meaningful: arguably the most basic mechanical consequence of locomotion is to change the body’s center of mass position; stepping variables are routinely measured in the laboratory; and our model’s control inputs, the push-off impulse and hip spring, represent net toe-off impulses and the action of hip abductors during leg swing, respectively. However, the model’s limited kinematics and actuation preclude it from being used, by itself, to study the range of mechanisms that may be used to maintain viable gait, which anchor in the human body the hierarchical control scheme identified in this study. Such mechanisms include those employing degrees of freedom at the ankles, hips, torso, or upper limbs [1, 54–58]. Furthermore, the three control inputs in our model were adjusted discretely at each step; however, under general conditions, particularly away from steady upright walking, one expects other generalized muscle forces to play a significant role, and for these forces to be modulated in continuous time. Indeed, the control inputs, viewed as being generated by individual, “lumped” actuators in our model, must ultimately be implemented by the human central nervous system [11, 12] at the level of muscles and reflex loops. Models incorporating spinal control and reflex pathways have been proposed [49, 59, 60] that provide a possible pathway toward anchoring the integrated template identified here in more realistic neuromechanical models.

We believe our study has broad relevance for at least two reasons. First, as discussed in [1], the literature to date overall supports the idea that foot placement is the dominant mechanism for maintaining gait stability, with other mechanisms being recruited as supplemental or compensatory strategies. Thus, our integrated template model, possibly with appropriate modifications to the task level regulation policy, may help interpret stepping dynamics observed in a variety of walking experiments (involving, e.g., lane changes [44] or other maneuvers). Second, the generalizability of our study stems from its articulation of an overarching, and experimentally testable, control architecture for goal-directed walking, together with an analysis of the geometrical dynamics that results from it. Even in much more realistic, “anchored” walking models, with many degrees of freedom and concomitantly higher-dimensional actuation schemes, steady gaits capable of satisfying specific task-level goals should, in general, be expected to lie on GEMs in the system’s extended state space. We further conjecture that at least a subset of the steady gaits contained in these GEMs will typically form semistable sets.

## Data access

The code for this study can be found in the Zenodo repository at https://zenodo.org/doi/10.5281/zenodo.12636582 [61].

## Acknowledgments

This work was supported by the U.S. National Institutes of Health grant No. R01-AG049735. The authors thank Christopher Rahn at Penn State for a helpful discussion on successive loop closure.

